# Deep computer vision reveals the mediation of thermal-melanism and body size by precipitation and sex in a threatened alpine butterfly, *Parnassius smintheus*

**DOI:** 10.1101/2024.02.05.578938

**Authors:** Vaughn Shirey, Rhea Goswami, Greg Latronica, Arshan Goudarzi, Naresh Neupane, Greg A. Breed, Leslie Ries

**Author notes:** **Funding:** VS was supported by a National Science Foundation Graduate Research Fellowship (#1937959) and a David H. Smith Conservation Research Fellowship while conducting and writing about this work. **Author Contributions:** VS came up with the original study design. VS, RG, GL, AG, and NN collected and annotated the training and validation data. VS and RG constructed the computer vision pipeline. VS conducted the statistical analysis. GAB and LR provided helpful guidance on the work. VS wrote the initial manuscript draft, and all authors provided feedback and edits. All authors approved the final manuscript for submission. **Note to Editor and Reviewers:** Significant portions of this work can be found in the online and print versions of the lead author’s doctoral dissertation and should not be considered plagiarism.

## Abstract

Insect morphologies are strongly tied to selective forces, yet due to variation in these forces and finite resources, insects must strategically invest in select morphologies while deprioritizing others. Melanism and body size may be one such potential trade-off that insects navigate and these two factors are important for fecundity, dispersal, thermoregulation, anti-desiccation, and immunity. In this work, we examined how sex and environmental factors mediate a potential body-size/melanism trade-off in the cold-adapted butterfly *Parnassius smintheus* (Lepidoptera: Papilionidae). We used deep computer vision approaches and museum specimen photography to process over 1,000 images of the species. We found that body-size and melanism are strongly mediated by temperature and elevation (thermal-melanism hypothesis) and that precipitation mediates these slopes for males and females differently. Notably, under the wettest conditions, females exhibit stronger concordance with the thermal-melanism hypothesis while the relationship for males is inverted, suggesting increased competition among males in cold-wet environments. Our results highlight the importance of considering sex when examining how the environment influences intraspecific morphological variation, especially under projected scenarios of global climate change.

## 1.0. INTRODUCTION

The variety of color has captivated the attention of artists and the general public throughout human history (Cuthill *et al*. 2017; Finlay 2007). Morphological color patterns are also a critical component of understanding ecology and evolution. As the most biodiverse group of animals described on the planet, insect color and morphology can evolve locally over relatively short timescales and in response to environmental conditions such as temperature, precipitation, and biotic interactions (Boggs *et al*. 2003; Clusella-Trullas & Nielsen 2020; Kingsolver 1988; Pegram *et al*. 2013). Insect morphology can also be plastic, responding to developmental conditions to produce varying phenotypes (Karl *et al*. 2009; Roland 1982). These factors have led to the proliferation of color diversity for various utilities among insects. For example, color is critically important for signaling conspecifics and butterflies are a model system for understanding color and its perception (Martin & Reed 2014; Rutowski *et al*. 2005). Additionally, insects are masters at using color and morphology to avoid predation via crypsis or aposematism (Nadeau *et al*. 2016; Vallin *et al*. 2006). Color also serves additional functional roles in insect physiology.

Within many insect species, there is often a high degree of variation in body and wing coloration patterns. As ectotherms, regulating body temperature is a critical component of insect metabolism and behavior and is strongly influenced by coloration (Trullas *et al*. 2007). Melanin, a class of important dark-colored pigments, is the best-studied form of coloration concerning insect thermoregulation (True 2003; Trullas *et al*. 2007). Since melanin is a dark pigment, it absorbs solar radiation converting visible wavelengths to heat (as opposed to white pigmentation which reflects this radiation). This thermal energy can be directed into an insect’s body, allowing for more efficient thermoregulation and warming of flight muscles and other tissues (Guppy 1986; Roland 1982, 2006). Higher rates of body/wing melanism have been reported from individuals that occur in colder habitats or at higher elevations – underscoring the likely functional importance of melanism in insect thermoregulation (Guppy 1986; Karl *et al*. 2009; Roland 1982, 2006; Trullas *et al*. 2007). In these environments, it is thought that investment in melanism confers increased basking efficiency and allows individuals to spend more time flying and less time engaging in thermoregulatory behaviors (Dufour *et al*. 2018; Guppy 1986; Roland 1982, 2006). Less time spent basking affords individuals larger time budget for resource and mate acquisition, increasing the fitness of more melanized individuals in colder climates. Past research has shown that melanistic patterns in response to temperature depend on species-specific climate sensitivity. For example, two genera of butterflies in the Andes exhibit opposite melanization patterns in response to an elevational gradient which is likely driven by differential responses to temperature and solar radiation (Dufour *et al*. 2018). Herein, we refer to this functional role of melanism as the thermal-melanism hypothesis (Figure 1a).

**Figure 1.**
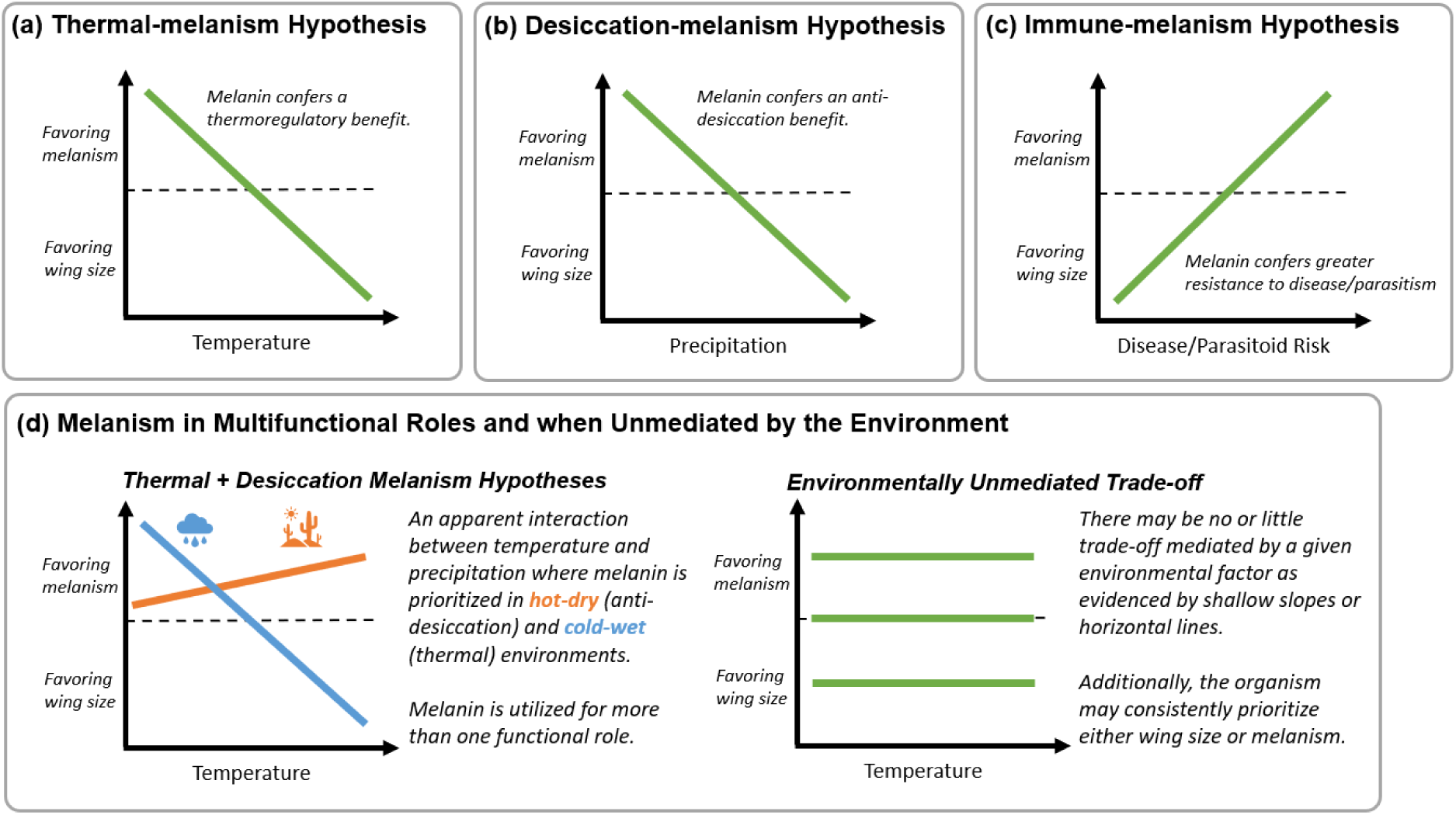
Our hypotheses surrounding the mediation of wing size and melanism along varying environmental gradients. Panel (a) illustrates the expectation under the thermal-melanism hypothesis with melanism being favored in low temperature regions. Panel (b) illustrates the desiccation-melanism hypothesis with melanism elevated in dry regions. Panel (c) illustrates the expectation under an immune-melanism hypothesis (not directly tested here). Finally, alternative scenarios may emerge due to combinations of selective pressures or the trade-off may be unmediated by our examined environmental axes.

Melanism, however, can serve several other important functions in addition to its thermoregulatory properties. The family of melanin molecules is hydrophobic (Nosanchuk & Casadevall 2003; Wu & Hong 2015), and in some insects it has been shown that higher melanin levels in the cuticle can confer desiccation resistance (King & Sinclair 2015; Krupp *et al*. 2020; Ramniwas *et al*. 2013). Here we refer to this hypothesis, as a desiccation-melanism hypothesis (Figure 1b). In some insects, such as tree wetas (Orthoptera: Anostostomatidae), higher melanic morphs have demonstrated the ability to reduce the rate of water loss by as much as 55% compared to non-melanic phenotypes (King & Sinclair 2015). Conversely, in laboratory experiments with *Drosophila* flies, no support was found for the role of melanism in promoting survival (although higher melanized individuals did experience water loss at a lower rate)(Rajpurohit *et al*. 2016). Like thermal-melanism, the importance of melanin in water conservation may be highly context and species-dependent.

Melanin also plays a central role in insect immunology. When faced with foreign body intrusion or disease, melanin is used by insects to isolate potential pathogens via encapsulation (Cotter *et al*. 2008; Dubovskiy *et al*. 2013; Gillespie and *et al*. 1997; Götz 1986; Kanost & Gorman 2008). Disease risk may be elevated in warmer, wetter regions, and individuals in such habitats may produce more melanin and thus appear darker in response to these immunological threats (Dobzhansky 1950; Janzen & Schoener 1968; Roslin *et al*. 2017). We refer to this phenomenon as the immune-melanism hypothesis (Figure 1c). The relationship between increased humidity and melanism has been recognized since 1833 (Gloger 1833), and this has become known as Gloger’s Rule. The underlying drivers of this rule vary, and studies that can support specific drivers of this pattern are rare (Burtt Jr & Ichida 2004; Jablonski & Chaplin 2010). Numerous other driving forces for Gloger’s Rule include increased selection pressure for camouflage and other forms of inter-and intraspecific interaction (Delhey 2019).

The potential multifunctionality of melanism in insects makes disentangling the environmental drivers of these pigments difficult. Further, melanin may be costly to synthesize, especially for herbivorous insects, since it requires exogenous tyrosine or phenylalanine (the latter an essential amino acid) to be acquired through feeding (Hearing 2011; Lemoine *et al*. 2020). Other research has confirmed the importance of host plant quality and stressors experienced during development as likely drivers of insect morphology and color quality (Pegram *et al*. 2013; Talloen *et al*. 2004). Since these amino acids are also important for growth, individuals may need to divert resources to growth or melanism. Such a size-melanism trade-off would mean that insects may need to optimize allocation of physiological resources towards either size or increased melanization.

Considering the potential functions of melanin in insects and the basic assumption that “bigger is better” for both males and females, we might expect that (a) increased melanism is prioritized over body size in cold regions (thermal-melanism, Figure 1a); (b) melanism is elevated over body size in dry regions (desiccation-melanism, Figure 1b); or that (c) melanism is elevated over body size in warm, wet regions (immune-melanism, Figure 4c). Further, such a trade-off may be complicated by combinations of the above environmental factors that might lead to more complex patterns. For example, if the thermal-melanism and desiccation-melanism hypotheses are both important, we would expect melanism to be elevated in both cold and hot-dry environments (Figure 1d). Under these circumstances, determining if there is an interaction between temperature and precipitation will be critical to elucidating underlying mechanisms.

All else being equal, larger individuals likely garner a fitness benefit, given the generally positive relationships between body size and fecundity in insects (Akman & Whitman 2008; Alcock 1994; Berger *et al*. 2008; Honěk 1993); however, the extent to which these hypothesized environmental conditions mediate trade-off is difficult to test experimentally across a species’ entire geographic range because designs of this nature are costly or fraught with logistical challenges. For example, common garden experiments, likely the most powerful approach to teasing apart drivers of melanism and body size, would be difficult to scale (Benito Garzón *et al*. 2019). Despite this challenge, large-scale studies comparing the morphology of specimens collected across their geographic range provide an opportunity to test this framework by linking each specimen to the historical climate in which it developed.

This approach relies on the variability in precipitation and temperature across space and time to disentangle evidence for priority investment in melanism or size as a quasi-experiment. Such an approach is increasingly possible because of the global effort to digitize billions of museum specimens and make those records publicly available on aggregating websites such as the Global Biodiversity Information Facility (GBIF) (Sanders *et al*. 2023). While this effort initially focused on transcribing label data to amass records of occurrences (species, location, date of collection, etc.), specimen photographs are now also being made available in online portals, providing a growing data source for exampling morphological variation across environmental gradients (Wilson *et al*. 2022). Nevertheless, studies using this more observational approach will be constrained by the number of available specimen photographs and the degree to which temperature and precipitation vary and covary across a species’ range. With these advancements, testing for potential size-melanism trade-offs is now possible at scale for some insect species through advances in deep computer vision, a form of artificial intelligence. There are many logistical challenges for leveraging these types of data; however, deep computer vision approaches can collect massive amounts of morphological data from specimen photographs in relatively short amounts of time compared to manual scoring (Høye *et al*. 2021; Wilson *et al*. 2022). Additionally, as new photographs become available, automating the extraction of morphological data will allow for future validation of hypotheses. In this work, we applied deep computer vision to extract a large body of morphological data for the alpine butterfly species, *Parnassius smintheus*, Doubleday, [1847] (Lepidoptera: Papilionidae), and examine potential size-melanism trade-offs across its North American range.

*P. smintheus* is an ideal study system to examine the potential trade-off between size and melanism for several reasons. First, *P. smintheus* is a cold-adapted species with a geographic range extending from northern New Mexico into the southern Yukon Territory (Figure 2). It is largely monophagous, feeding mostly on the stonecrop species *Sedum lanceolatum* Torr. (Saxifragales: Crassulaceae) (Nishida & Rothschild 1995) and therefore we can reduce the complexity of having to consider the resource quality of different host plant species in a downstream analysis. Upon mating, males impart a sphragis, or copulatory plug, on the females that prevents further copulation (Matter et al. 2012) and so the mating system is fairly constrained to pairwise interactions. *P. smintheus* is a typically alpine species residing in open-canopy habitats. However, it can also be found at lower elevations in steppe habitats, especially in the northern periphery of its range (based on collective field experience in the Yukon). Notably, individuals of this species are reluctant to traverse closed-canopy habitats, which likely contributes to a strong spatial structure of localized populations (Matter & Roland 2002; Matter et al. 2004; Ross et al. 2005). Flight behavior is associated with sex, and males of this species are typically more aerially active than females – aggressively patrolling habitats in search of females to mate with and assessing potential habitat quality (Matter & Roland 2002). However, females are more likely to traverse closed canopy habitats and disperse over greater distances (Goff *et al*. 2019).

Like other montane species, global climate change is forecasted to be a major threat to the survival of this species. Threats induced by climate change can include the direct impacts of rising temperatures and shifting precipitation regimes on the species. For example, Roland and Matter 2016 found that extreme early winter weather patterns are associated with low adult abundance in the following flight season (Roland & Matter 2016). Additionally, tree-line encroachment into alpine meadows may reduce viable habitat for the species and connectivity between patches (Ross et al. 2005; Matter et al. 2020). Despite this, Matter and others indicate that neither climate nor habitat loss are reliable predictors of population size or metapopulation network stability and that interannual variation in dispersal capacity for this species can be quite high (Matter & Roland 2017; Matter et al. 2020). Characterizing the variation in morphology for the species may explain some of the species’ ability to mitigate threats through local adaptation.

*P. smintheus* exhibits remarkable morphological variation across its range, making it a particularly excellent focus for this study. The species is sexually dimorphic (Figure 2a), with females being redder, darker, and more transparent than males. Melanin is considered critically important for flight behavior in the species, especially when located proximal to the flight musculature (Guppy 1986). Elevated wing melanism likely contributes to more efficient basking behavior and, consequently, increased flight capacity in cooler temperatures (following the thermal-melanism hypothesis). While the field studies conducted by Guppy in the 1980s has provided evidence to support the thermal-melanism hypothesis in *Parnassius*, notable questions remain with respect to how melanin might be optimized across the range, and how this may vary by sex and wing. For example, in 1989, Guppy found little evidence that melanism in female *P. phoebus* (under which *P. smintheus* has long been considered a subspecies) was driven by environmental conditions, although sample sizes were small and localized (Guppy 1989). At larger spatial scales, a study in the congener species, *P. clodius*, found that solar radiation exposure strongly predicted melanin following the thermal-melanism hypothesis, with individuals exposed to higher solar radiation expressing less wing melanism (Zaman et al. 2019). To our knowledge, no study has examined trade-offs between wing size and melanism in *P. smintheus.* Given the importance of temperature in this species’ life cycle and potential climate-oriented threats, and the physiological cost of producing melanin, we explore that potential here. Specifically, we aimed to accomplish the following key goals:

1. Evaluate evidence for a size-melanism trade-off in *P. smintheus* across its range using a historical dataset of specimen photographs.
2. Examine this potential trade-off along key environmental axes (temperature, precipitation, and local elevation), and interactions among them, experienced by each individual during their developmental life stages.
3. Examine different patterns in these trade-offs by sex and between fore-and hindwings with particular focus on hindwings which have been previously indicated to be important for thermal regulation.

## 2.0. METHODOLOGY

### 2.1. Photography and Deep Computer Vision Pipeline

All data for this analysis were extracted from photographs of museum specimens that included the location (either a coordinate location or description of a locality to be converted to a coordinate) and year of collection. All photographs met a set of minimum criteria for inclusion, namely that they contained some sense of scale (either a scale bar or ruler) and a color target card so that photographs could be color-corrected for fair comparison. These two items in the photo ensure that unit-based (e.g., centimeters) as opposed to pixel-based measurements could be calculated and that color was assessed without bias across different camera and lighting setups. Standardization of museum specimen photographs provides the opportunity to extract morphological information and tie such data to standard occurrence information, such as when and where the specimen was collected.

We took two approaches to aggregate photographs of *P. smintheus* specimens. First, all available specimen image data were downloaded with corresponding occurrence information from the Global Biodiversity Information Facility (GBIF) (GBIF 2022), Integrated Digitized Biocollections (iDigBio), and the Symbiota Collections of Arthropods Network (SCAN) (Heinrich et al. 2015). Since GBIF, iDigBio, and SCAN often aggregate the same specimen data, all duplicate records were removed from our analysis. We note that most of the photographs from these databases are from the Yale Peabody Museum, which used a Canon EOS T3 with a Canon 60mm macro prime lens to take the majority of the photos (Dr. Lawrence Gall, personal communication) and from the LepNet digitization project (Seltmann et al. 2017). Second, we visited three natural history museums: The American Museum of Natural History (New York, New York); The Yale Peabody Museum (New Haven, Connecticut); and The McGuire Center for Lepidoptera and Biodiversity Research – Florida Museum of Natural History (Gainesville, Florida), to add additional specimen photographs to our dataset. We used either a Canon EOS 7D with 24055 Canon Zoom lens (Dr. Andrei Sourakov; personal communication) or a Nikon D7500 DLSR camera with Yongnuo 50mm prime lens to photograph dorsal images of butterfly specimens during these visits. In total, we amassed 1,756 images with associated occurrence data.

From each specimen, we extracted the latitude and longitude of the capture location and the year of the specimen’s collection. If these data were unavailable, we would use a latitude or longitude coordinate using the verbatim description of the collection locality on the specimen label. We used GeoLocate (https://geo-locate.org/) to perform this historical georeferencing. Specimens that lacked this information or could not be resolved to a latitude and longitude point were excluded from the analysis. When sex was not already provided with the occurrence data, a determination of each specimen’s sex was made visually. In the cases where this was ambiguous, these records were excluded from downstream analysis (*n = 1*). These processes resulted in 1,470 images for our analysis (Figure 2b) (1,379 came from existing online portals and 91 from our collection visits of specimens). Of these, 1,063 were males and 407 were females, with ratios of each roughly even across years, although captures were slightly male-biased at the beginning and female-biased at the end (Figure 2c). From each photo, our goal was to outline the boundary of all four wings and calculate the wing area and extent of melanism apparent in each. Rather than annotate our photographs manually, we opted to use deep computer vision to automate this process. This allowed us to process our specimen photos more consistently and efficiently and also provided a mechanism to scale future efforts as the number of photographs of museum specimens continues to grow (Wilson et al. 2022b).

**Figure 2.**
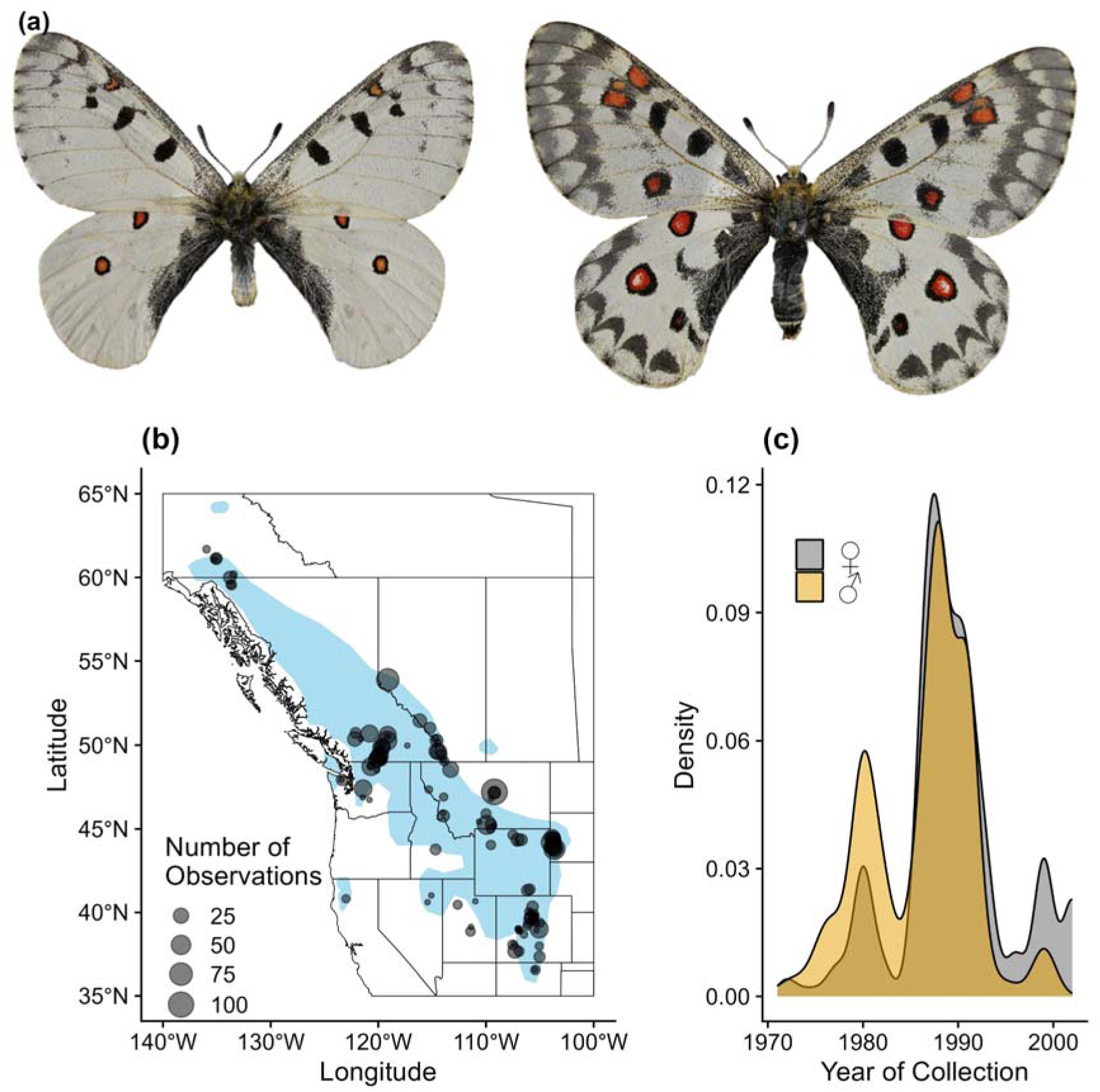
Examples of (a) sexual dimorphism in P. smintheus (male left, female right); (b) a map showing the geographic distribution of photographic records used in the analysis overlaid on the species range (in blue shading); and (c) a density plot showing the temporal distribution of records used in the analysis. Males are represented in gold, while females are represented in grey.

Deep computer vision has been heralded for its potential to exponentially increase our ability to extract morphological data from natural history museum collections (Høye et al. 2021; Wilson et al. 2022b), and thus allow us to better tap into the information available from individual specimens across large spatial and temporal extents. We detail the construction of our deep computer vision pipeline in Supplemental Material S1, but briefly, key features for analysis were extracted from image files in three main steps. First, we used the color target card in each image to adjust the photograph to make color metrics directly comparable among all records using ImageJ. For the next two steps, we used convolutional neural networks (CNNs), a powerful tool for pattern recognition (Høye et al. 2021; Wilson et al. 2022b). We used one of these CNNs to detect the scale bar in each image and save the conversion factor from pixels to centimeters for that image. We refer to this neural network as “ScaleNet.” At this point, we obtained the necessary information to compare wing color and size across specimens. The next step involved extracting the metrics used in our analysis from the wings. We trained a second CNN, which we refer to as “WingNet,” to segment each wing and extract from these wings the total wing area, total melanized area, and the proportion of the wing area that was melanized. Details on these neural networks can be found in Appendix A (Supplemental Material S2). To perform this work, we relied heavily on transfer learning from Meta’s Detection2 library (Wu *et al*. 2019).

After segmenting each wing, we used a binary thresholding procedure to define each pixel as either melanized or not. In a L*a*b* color space, we specified that melanized pixels must have a luminance value (L*) below 10 and that the a* and b* values fall between -10 and 10 (this was to prevent colors like dark red, orange, blue, or green being included as melanism). A L*a*b* color space is a way of defining color that is close to human perception (Schanda 2007). All colors falling outside of this threshold were not counted as melanized. Since one of the key assumptions of our analyses and hypotheses is that there is a trade-off between size and melanism, we first examined our wing metrics to establish that this relationship is evident in *P. smintheus*. As expected, results show that wing size and melanism have a significant, negative correlation (Figure 3), allowing our overall larger analysis to proceed.

**Figure 3.**
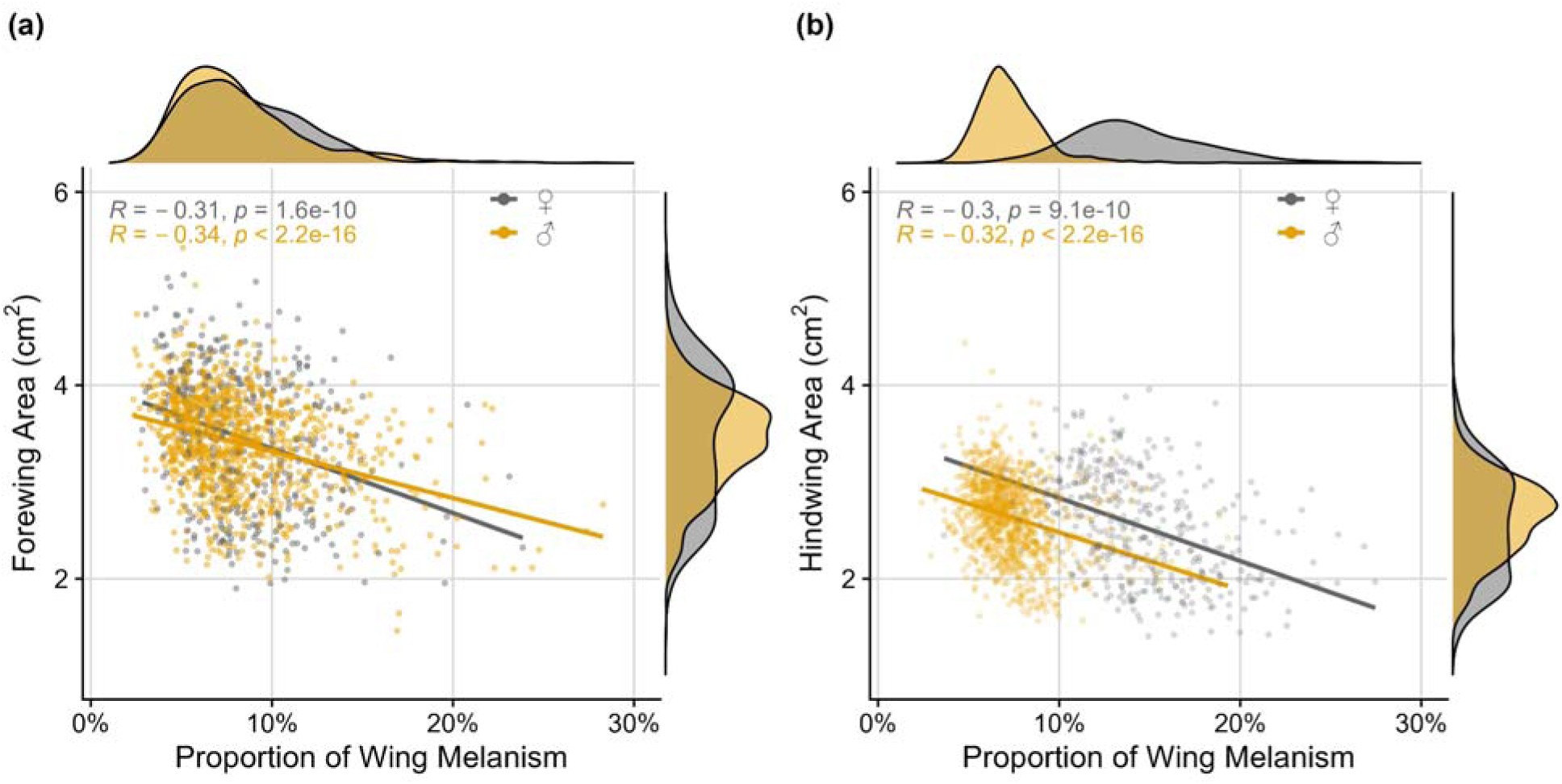
Summary distributions of wing area and the proportion of melanism in (a) forewings and (b) hindwings. Lines represent Pearson’s correlation analysis results between these two axes. A significant negative correlation exists between wing area and melanism for both sexes and in both wings. Males are represented in gold, while females are represented in grey.

### 2.2. Analysis of the Size-melanism Trade-off

Our response variable for all analyses was a trade-off metric. We calculated this metric as the difference in z-scores between the proportion of melanin on the wing and wing area. This metric is positive when individuals invest more heavily in melanism and negative where they prioritize investment in wing area (e.g., □□□□□ - □□□ □□□□□□ = % □□□□□□□□ *Z-score- □□□□ □□□□ Z-score*). Using this trade-off metric, we examined how investment in size versus melanism may be mediated along the axes of environmental gradients, specifically temperature, precipitation, and elevation.

To estimate the conditions that each individual may have experienced, we assume they went developed in roughly the same region in which they were captured (roughly September through May), and calculated the average temperature and precipitation amount for each individual’s capture location. In other words, if an adult specimen was collected in 1980, we inferred the average developmental temperature/precipitation from September 1979 to May 1980, or when the individual was an egg, caterpillar, and pupa. We acquired these climate data at a monthly, 1-degree spatial resolution from the National Ocean and Atmospheric Administration’s 20th Century Reanalysis project (Compo et al. 2011; Slivinski et al. 2019). As alpine species, elevation may also be important and, in our data, may represent climatic conditions at smaller spatial scales (5-minutes) than the climate data (1-degree). Elevation was extracted from a digital elevation model, ETOPO5 (National Geophysical Data Center 1993). We determined the degree of correlation between temperature and precipitation and found that they were not strongly correlated across our study region (Pearson’s R = -0.2). Based on our framework for observing trade-offs (Figure 1), existing variability in temperature and precipitation are structured in a way that allows the thermal-melanism and desiccation-melanism hypotheses to be more directly evaluated.

We constructed a series of spatiotemporal mixed models to assess the relationship between environmental predictors and trade-offs evident in wing morphology. To account for non independence among our samples, we included a random-intercept term based on location and year in our models to capture spatiotemporal covariance structure. We assumed that the relatedness between observations followed a linear decay function through time and an exponential decay function across space (the strength of this exponential decay was varied across iterations of the model to examine the strength of this spatial covariance structure). We used the Akaike Information Criterion (AIC) (Akaike 1998) and log-likelihood calculations to select the most performant model from this process. Details on the construction of this covariance structure and varying spatial dependence are provided in Supplemental Material S2. The general form model we used to investigate the trade-off was defined by Equation 1.

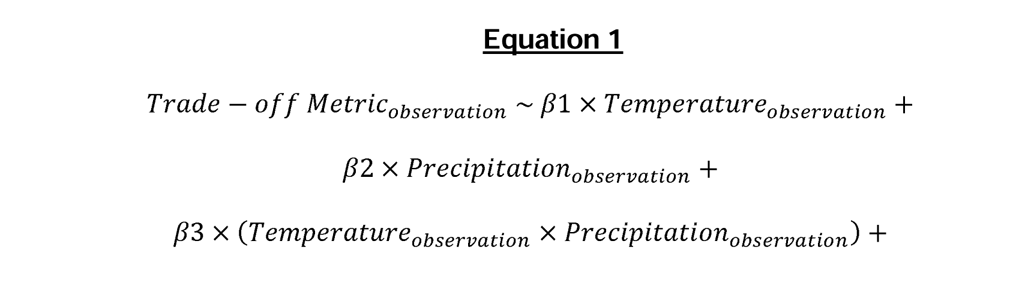

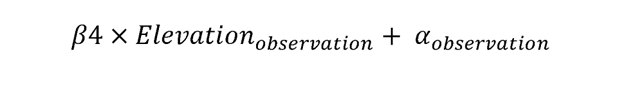

where *{*1 indicates the effect of inferred developmental temperature on the trade-off. *{*31 indicates the effect of inferred developmental temperature on the trade-off, {32 indicates the effect of inferred developmental precipitation on the trade-off, {33 indicates the interaction effect of temperature and precipitation on the trade-off, {34 indicates the effect of elevation on the trade-off, and a_observation_ refers to the intercept of a particular observation expected to follow the spatiotemporal covariance structure specified above so that observations closer in space and/or time will be more similar to each other than distant observations. To assess whether multicollinearity was an issue in our analysis, we calculated variance inflation factors (VIFs). Generally, VIFs greater than ten are considered to cause problems in regression analyses (although see O’brien 2007 for how thresholds may be controversial in practice) and indicate that correlated variables should be dropped from the analyses. We ran a separate model for both sexes and wings. All analyses were conducted in R v. 4.2.2 (R Core Team 2022) with the package “spaMM” (Rousset & Ferdy 2014). The code used in this analysis is available on GitHub at XXX and in a static Zenodo repository at XXX (DOI XXX).

### 3.0. RESULTS

We found a negative correlation between wing size and the proportion of wing melanism (Figure 3, but see SI for further details on this relationship). *P. smintheus* females exhibited greater variation in both forewing and hindwing area compared to males (Figure 4). Females were also more variable in hindwing melanism, and were generally darker than males. Absolute melanized and total wing areas were positively correlated on both wings, but this correlation is notably stronger in hindwings (Supplemental Figure S3). Generally, as wing area increases, so does the absolute area of wing melanism; however, we note that this correlation is not perfect, and there is variation in this pattern for both forewings and hindwings. The total melanized wing area and the proportion of the wing melanized were strongly positively correlated on both wings, with greater variation around this correlation in the hindwings, especially for females (Supplemental Figure S4).

**Figure 4.**
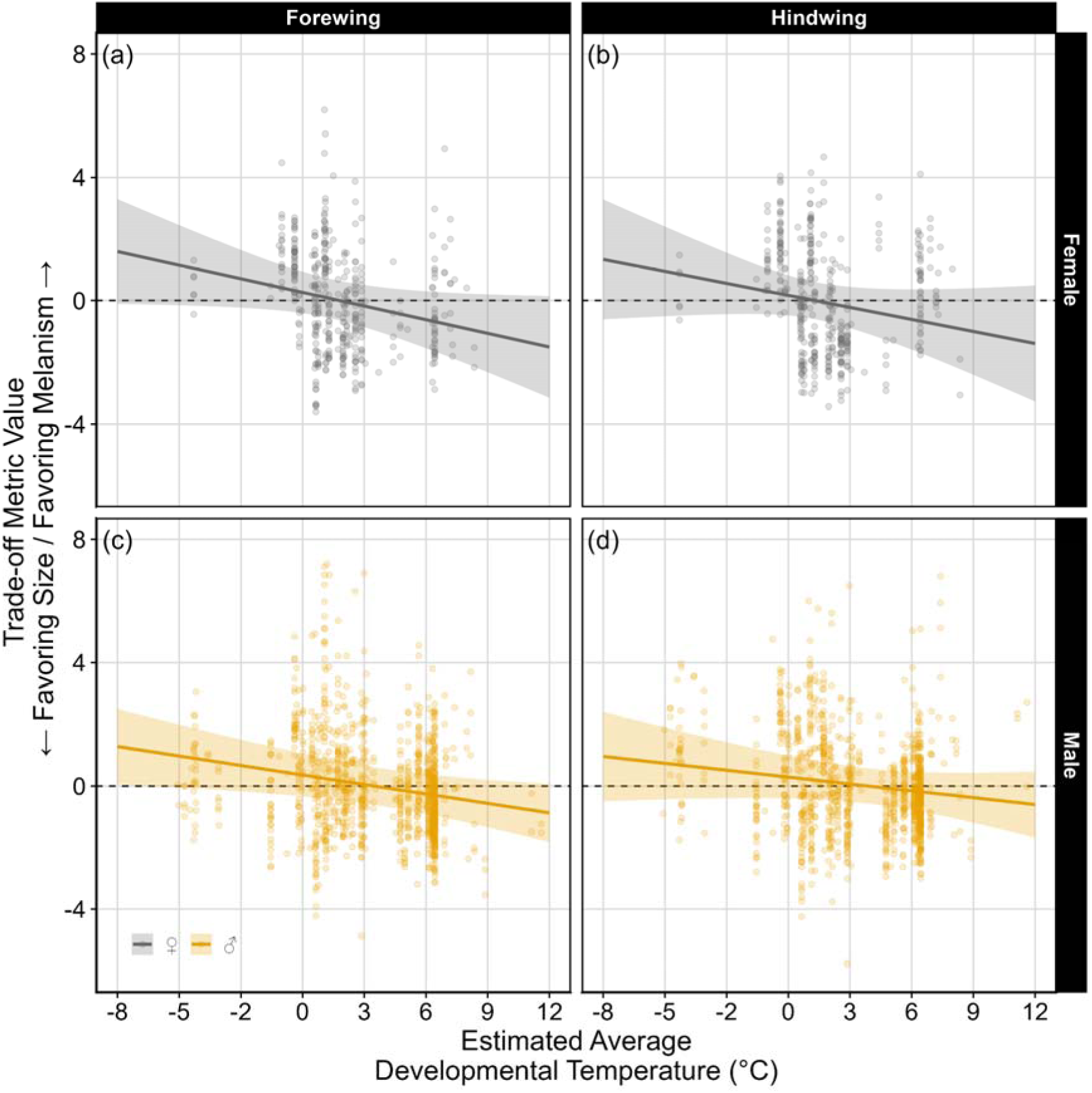
Partially-dependent effects plot of inferred developmental average temperature on (a) female forewing, (b) female hindwing, (c) male forewing, and (d) male hindwing size-melanism trade-offs. Positive values indicate favoring melanism, while negative values indicate favoring size. The relationships between temperature and the trade-off metric are significant for female and male forewings (a, c) but not the hindwings (see Table 1 for model statistics).

Our *a priori* hypotheses regarding the potential trade-off between wing size and melanism in *P. smintheus* garnered varying support with notable differences across sexes and wings. The thermal-melanism hypothesis gained the most support from our models (Table 1, 2, Figure 4, 5). Generally, butterflies favor melanin when they experience lower developmental temperatures; however, this relationship was only significant for male and female forewings (Figure 4a,c). In contrast, at more local scales, individuals at higher elevations prioritized melanism over wing area across both sexes and wings (Figure 4). These results are largely concordant with our initial hypotheses presented in Figure 1a. The desiccation-melanism hypothesis was not supported by our models, as evidenced by largely low parameter estimates and high confidence intervals (Table 1, 2, and Figure 6, Supplemental Figure S5).

**Figure 5.**
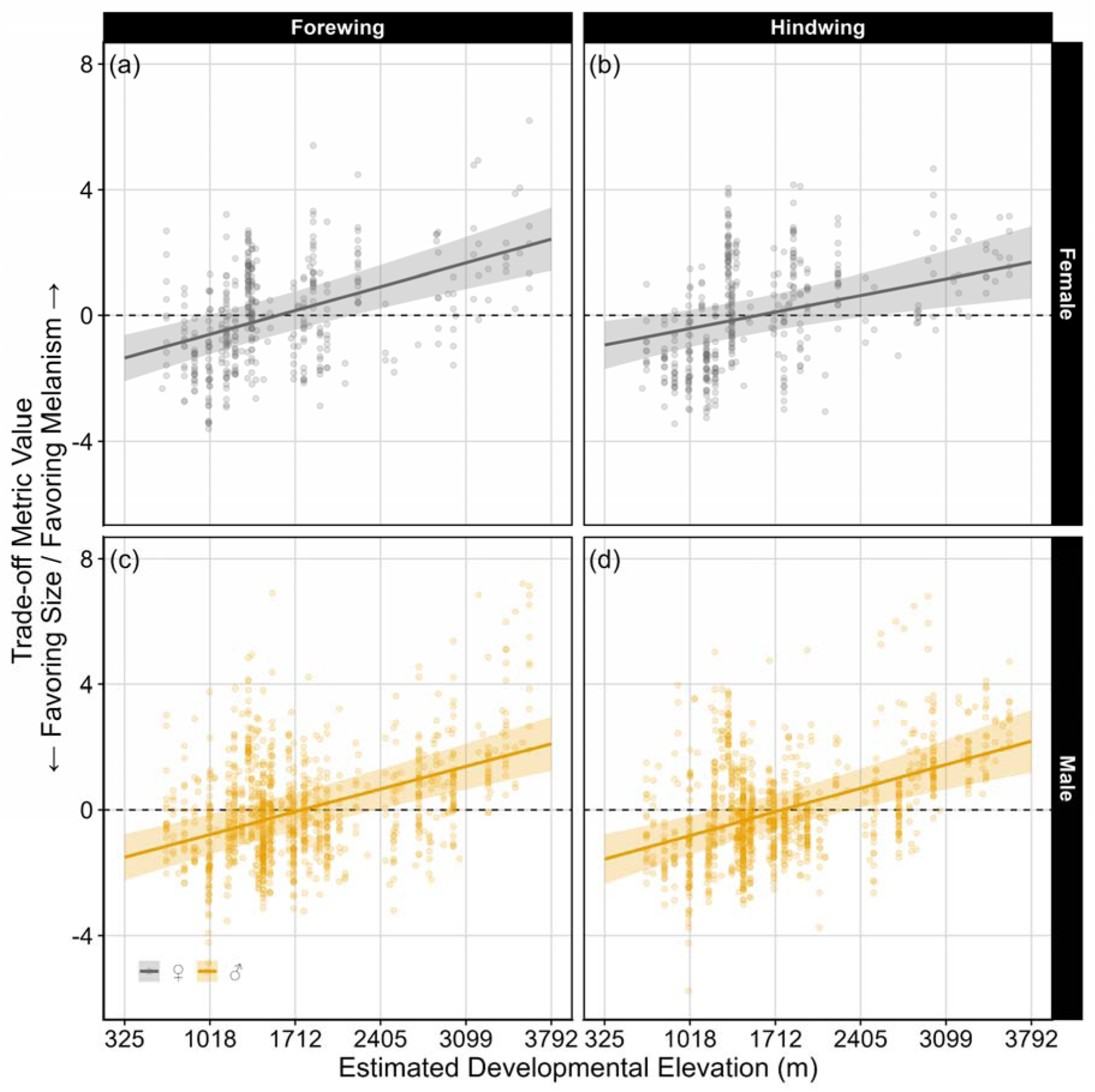
Partially-dependent effects plot of inferred developmental elevation on (a) female forewing, (b) female hindwing, (c) male forewing, and (d) male hindwing size-melanism trade-offs. Positive values indicate favoring melanism, while negative values indicate favoring size. The relationships between elevation and the trade-off metric are significant for all sexes and wings (see Table 1 for model statistics).

**Figure 6.**
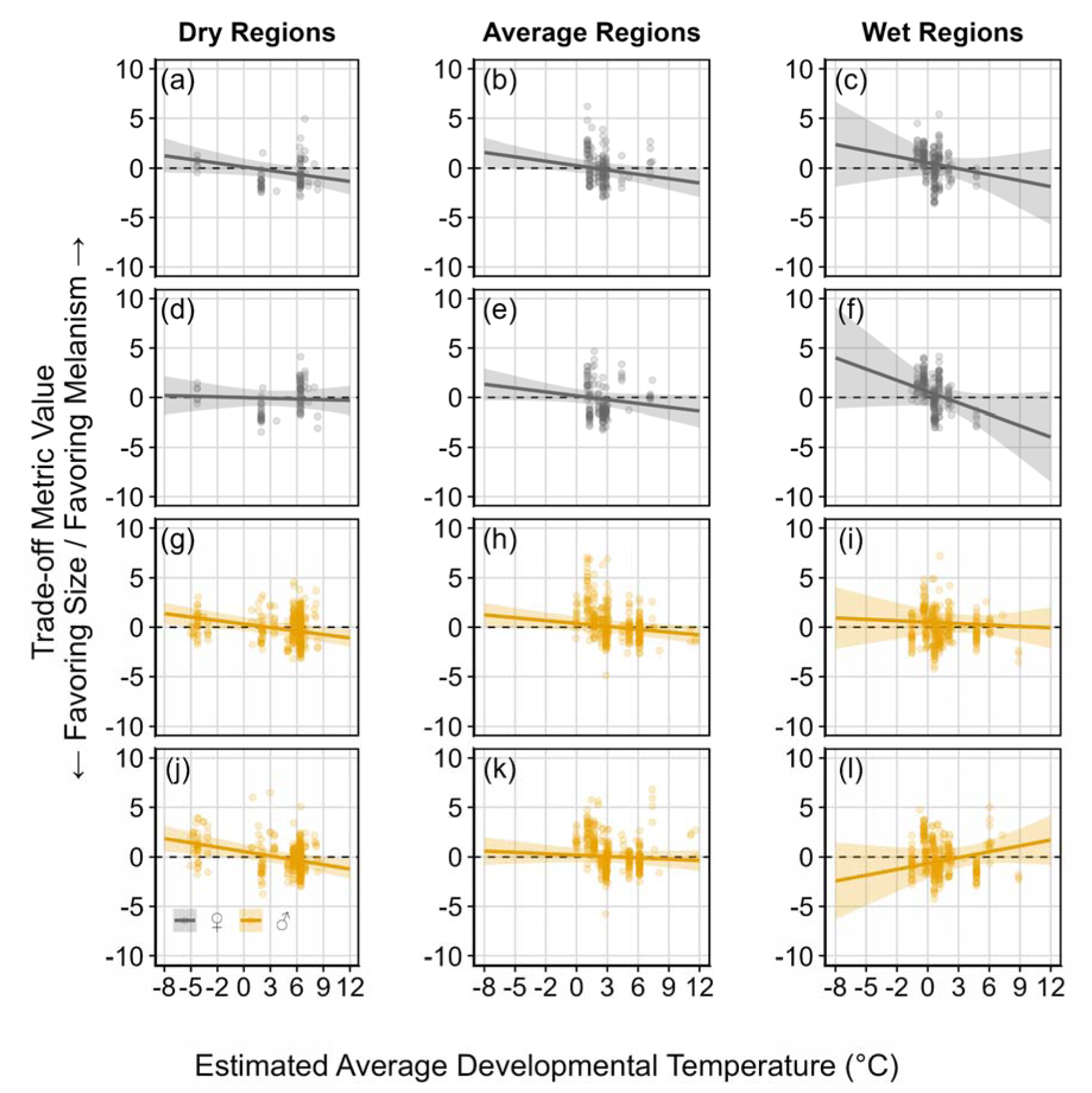
Partially-dependent effect plots of the interaction between inferred developmental precipitation and the average temperature on (a-c) female forewing, (d-f) female hindwing, (g-i) male forewing, and (j-l) male hindwing size-melanism trade-offs. Positive values indicate favoring melanism, while negative values indicate favoring size. The interaction effect between precipitation and temperature is significant for male hindwings (j-l) only (see Table 1 for model statistics).

**Table 1.** Parameter estimates for the top model when predicting the forewing size-melanism trade-off metric in both sexes. Asterisks denote significant predictors of the trade-off at p < 0.05 (*) and p < 0.01 (**).

| Model Parameter | Estimate<br>[+/- SE]<br>(♀/♂) | Variance Partition<br>Coefficient<br>(♀/♂) | Pseudo-R<sup>2</sup><br>Statistic<br>(♀/♂) |
| --- | --- | --- | --- |
| Intercept | -0.17 [+/- 0.16]<br>-0.09 [+/- 0.12] | 0.32<br>0.34 | 0.44<br>0.34 |
| Temperature | -0.43 [+/- 0.17] **<br>-0.30 [+/- 0.11] ** |  |  |
| Precipitation | 0.05 [+/- 0.15]<br>0.11 [+/- 0.12] |  |  |
| Temperature x<br>Precipitation | -0.07 [+/- 0.12]<br>0.06 [+/- 0.11] |  |  |
| Local<br>Elevation | 0.76 [+/- 0.12] **<br>0.72 [+/- 0.11] ** |  |  |

**Table 2.** Parameter estimates for the top model when predicting the hindwing size-melanism trade-off metric in both sexes (note that a top model for each sex was selected). Asterisks denote significant predictors of the trade-off at p < 0.05 (*) and p < 0.01 (**).

| Wing and Sex | Model Parameter | Estimate<br>[+/- SE] | Variance<br>Partition<br>Coefficient | Pseudo-R<sup>2</sup><br>Statistic |
| --- | --- | --- | --- | --- |
| Female<br>Hindwings | Intercept | -0.07 [+/- 0.22] | 0.52 | 0.61 |
|  | Temperature | -0.32 [+/- 0.19] |  |  |
|  | Precipitation | -0.13 [+/- 0.18] |  |  |
|  | Temperature x<br>Precipitation | -0.30 [+/- 0.22] |  |  |
|  | Local Elevation | 0.53 [+/- 0.16] ** |  |  |
| Male<br>Hindwings | Intercept | 0.34 [+/- 0.17] * | 0.64 | 0.53 |
|  | Temperature | -0.20 [+/- 0.14] |  |  |
|  | Precipitation | -0.05 [+/- 0.15] |  |  |
|  | Temperature x<br>Precipitation | 0.29 [+/- 0.13] * |  |  |
|  | Local Elevation | 0.75 [+/- 0.14] ** |  |  |

Finally, we found precipitation strongly modifies the strength of the relationship between temperature and melanization (Figure 6). Higher degrees of melanism appears to be favored in both fore-and hindwings in both males and female when conditions are cold under dry to average precipitation regimes, yet the slope of this relationship is amplified in wet habitats for females (although the interaction was only statistically significant in male hindwings, the trend was present). Notably, there is an inverse relationship for male hindwings, where males appear to favor size in exceptionally cold-wet environments and melanism when those same moist environments are warmer (Figure 6l).

## 4.0. DISCUSSION

*P. smintheus* exhibits notable morphological variation consistent with a trade-off between investment in melanism and growth of larger wings (Figure 3). We tested several alternative hypotheses regarding the environmental gradients that potentially drive this trade-off based on over 1,000 images across their range and spanning 30 years, 1972-2002 (Figure 2c). Morphological responses to temperature were most consistent with the thermal-melanism hypothesis, among the three main hypotheses we tested (Figure 1).

Individual wing area was negatively correlated with the proportion of the wing melanized (Figure 3). Notably, this correlation is statistically significant even though the total area of the wing melanism scales positively with wing area (Supplemental Figure S3, S4, but note the relationship is not one-to-one and significant variation exists around this correlation). These associations support the size-melanism trade-off in *P. smintheus* and may suggest that individuals invest differently in producing either larger wings or more melanin. Similar support for the size-melanism trade-off has been found in insects (Safranek & Riddiford 1975; Ma et al. 2008; Dubovskiy et al. 2013); however, some studies suggest this trade-off may not be a “rule” but rather taxon or even environment-specific (True 2003; Stoehr 2006). Despite this, such a trade-off in *P. smintheus* makes sense from a resource-metabolic perspective since melanin and growth require nutrient acquisition from host plant consumption (Roff & Fairbairn 2013; Ethier et al. 2015). Further, melanin and body size are important traits for *P. smintheus,* given its specialization to colder habitats and sex-specific attributes such as intense male-male competition for mates.

The size-melanism trade-off was strongly supported under the thermal-melanism hypothesis. Most apparently, shifts towards prioritizing melanism over wing size were noted across wings and sexes with increasing elevation (Figure 5b), confirming that melanism is an important morphological feature at high altitudes (Guppy 1986). We found that, at more regional scales, inferred developmental temperature did not significantly predict the size-melanism trade-off except in male and female forewings (Figure 5a, c). Despite not being statistically significant, the hindwings exhibit the same directionality in slope (Figure 5b, d). Shallow estimates of the effect of a given environmental factor on the size-melanism trade-off metric may indicate that the morphology of the given wing is either highly conserved or not strongly mediated along that environmental axis or spatial scale. Microclimatic conditions experienced by lepidopteran larvae can strongly influence melanism and size (Kingsolver & Moffat 1982; Xing et al. 2016), and our findings suggest that this might indeed be the case for *P. smintheus*. We note that the positive result for the forewings was unexpected as most thermoregulation is thought to happen through the proximal melanism patches on the dorsal hindwings (Guppy 1986). We did not partition localized patches of melanism on our wings into other putative functional roles, but note that red ornamentation on *P. smintheus* is often highlighted by a highly melanized border (Figure 2a). If red ornamentation and melanism on the forewing covary together, one might expect additional, likely signaling, roles of ornamentation adjacent melanism to emerge. A recent analysis of *P. clodius*, demonstrated that bird predation risk did not strongly predict this red ornamentation (Zaman *et al*. 2019). However, one might expect predation risk to be elevated in hotter regions (Roslin *et al*. 2017), so further investigation of the role of redness and melanism on the forewing is needed.

Our analysis did not support the desiccation-melanism hypothesis. We found shallow slope estimates when examining the partial effects of inferred developmental precipitation on wing melanism and body size (Supplemental Figure S5). This is not to say that precipitation is not important, however. In our analysis of the interaction between temperature and precipitation, we found that precipitation strongly modifies the relationships between temperature and the size-melanism trade-off (Figure 6). The effect of temperature appears amplified in wet regions (Figure 6c, f) and is reversed for male hindwings (Figure 6l). Female investment in melanism over size in particularly cold-wet environments supports thermal-melanism as a driving mechanism behind this species’ morphology.

The case of male hindwings exhibiting an opposite pattern in cold-wet environments was more perplexing. We propose a few mechanisms for why males might favor larger hindwing sizes in cold-wet environments than melanism. First, male-male competition for *P. smintheus* for females can be quite intense, and large size may be a considerable advantage. Under the assumption that exceptionally cold-wet regions exhibit shorter growing seasons and shorter daily flight windows, the intensity of competition might be amplified within the brief phenological windows (Baena & Macías-Ordóñez 2015; Briones *et al*. 1998; Kinoshita 1998). Body size is understood to be a contributing factor to male mating success, especially when male-to-male competition is high in a variety of other organisms (Alcock 1994; Hagelin 2002). This hypothesis is complicated by untested trade-offs in our data - namely, a trade-off between emergence date, body size, and melanization and the need for detailed behavioral observation in the field (Zonneveld 1996).

A post-hoc analysis of the variance explained by the spatiotemporally structured random intercepts in our model confirmed some degree of spatial structure in our study system (Table 1, 2). Such variation may be explained by genetic relatedness between individuals but also by other local-scale factors such as host plant quality. Decomposing this variation into components explained by genetic relatedness versus other local-scale environmental conditions is needed; however, it is not currently possible with purely observational data. Despite this, attempts to examine the spatial scale at which additional variation might be best explained in our model is possible since our modulation of spatial dependence in this analysis may provide clues on where to look for explaining at what spatial scale additionally important environmental or genetic factors might be situated. For example, strong spatial relatedness among observations may be related to more local scale environmental factors like microclimate conditions and/or genetic relatedness structured by strong isolation-by-distance. In our analysis, additional variation in the model for forewings is situated at fine spatial scales (tuning parameter = 0.25), whereas in hindwings, these spatial scales differ by sex (tuning parameter = 1 for males and 2 for females). Understanding what these tuning parameters mean in a biological context is the next challenge for this research and will likely require additional investigation using covariates across spatial scales. Further, phenotypic plasticity in response to the environment and genetic determination of phenotype are not mutually exclusive; however, assessing how genotype might constrain plasticity requires formal experimental designs. This disentanglement can be supported with additional analyses using a sequence-based phylogeny and experimental common garden and inheritance trials. In such experimental designs, the heritability of melanism versus the plastic response of melanism to simulated temperature and moisture conditions, which aim to isolate other confounding effects, should be investigated using individuals sourced from across the species’ range.

We examined colors within the human-visible spectra in this analysis; however, ultraviolet reflectance has been shown to occur in *P. smintheus* to varying degrees (Shirey et al. 2021b). It is important to consider the mechanisms by which extrasensory (from a human context) color spectra can appear in species. For example, ultraviolet reflectance in other butterflies is often associated with mate finding (Silberglied & Taylor 1978; Robertson & Monteiro 2005; Rutowski et al. 2005). Anecdotally, from personal experience and other naturalists, *P. smintheus* females are typically mated shortly after the pupal life stage (thought to be driven by pheromone signaling), and thus the importance of ultraviolet reflectance in the species may not be as straightforward as mate finding strategy (McCorkle & Hammond 1985 for sister species *P. clodius*; Matter et al. 2012). Additionally, ultraviolet reflectance in *P. smintheus* appears co-located with red ornamentation on the wings (Shirey et al. 2021b), which may be related to warning coloration. However, as previously mentioned, Zaman et al. 2019 found that the relative abundance of insectivorous birds did not strongly predict red coloration in *P. clodius* (Zaman et al. 2019). Ultraviolet reflectance and red coloration on the wings may still suggest an important signaling component, but whether for predator avoidance or conspecific signaling remains to be tested. Behavioral observation and experimentation will be needed to associate ultraviolet reflectance with these hypotheses, while analyses similar to ours with broader spectra photography may be able to determine environmental determinants of this coloration and if any trade-offs exist between ultraviolet ornamentation and other morphological characteristics.

We also recognize that notable sampling biases exist in natural history collection data, even for butterflies, the best-sampled insects worldwide (Girardello et al. 2019; Shirey et al. 2021a). Due to this, we may not have aggregated a dataset in which the full range of habitable temperatures or precipitation regimes for *P. smintheus* were represented. This may hamper our inference, especially if sampling is sparse at climatic extremes, and thus, some points may have greater statistical leverage (e.g., extremely dry, wet, cold, or hot conditions). As digitization continues on the estimated billions of museum specimens, these analyses will not only have larger sample sizes but also, by examining patterns over large temporal spans, it will be possible to determine how these strategies can keep pace with climate change. Examining how ubiquitous the patterns discussed (and to what direction and magnitude they exist) across butterfly species may also point to greater evidence for the size-melanism trade-off hypothesis. Finally, other life-history conditioned trade-offs between size and melanin likely exist; however, the nature of opportunistic sampling in our dataset made testing for these trade-offs difficult. For example, there may be a trade-off between larger size and earlier adult eclosion date for protandrous males (Zonneveld 1996). Investigating a slew of potential trade-off mechanisms using deep computer vision approaches and natural history collections highlights the power of this approach. Indeed, computer vision has been heralded as potentially transformative for ecology and evolution (Høye et al. 2021; Wilson et al. 2022b). We enthusiastically support the continued digitization and photography of specimens to understand better how environmental factors might mediate differential investment of resources across the Tree of Life.

## Acknowledgements

Special thanks to Larry Gall, David Grimaldi, Akito Kawahara, Raymond Pudepis, Suzanne Rab Green, Andrei Sourakov, and Andrew Warren for access to various collections during the course of this work. We would like to thank Gina Wimp and Martha Weiss for helpful feedback on the initial draft of this work.

## SUPPLEMENTAL MATERIAL

### Supplemental Material S1: Computer Vision Pipeline

Since manual annotation of our complete image library would be a time-consuming process, we opted to train a series of convolutional neural networks (CNNs) to automate various image processing and feature extraction tasks needed to acquire color and other morphological information each image. Computer vision has been heralded as an important advancement in ecology and evolution, especially in entomology, as it can facilitate the rapid collection of information about species occurrence and morphology at scale (Høye *et al*. 2021). Our computer vision pipeline consisted of two CNNs: (a) ScaleNet, which determines the pixel-to-centimeter conversion for a given image, and (b) WingNet, which masks wings from the image for downstream size and melanism calculations. Before processing by ScaleNet or WingNet, each image was color-calibrated using the color-calibration chip in the image and the IJP-Color plugin for ImageJ. We used an L*a*b* colorspace for all images.

ScaleNet was used to detect scales/rulers within each image. ScaleNet used a training set of 70 images (validation set, 10 images), training over 2,000 epochs with a base learning rate of 0.025. After detection, ScaleNet outputs the pixel-to-centimeter conversion factor. Scale conversion factors were saved to a file to calculate the measurement of wing area and melanism. ScaleNet used transfer learning through the Detectron2 library with a Faster-RCNN-50FPN 3x backbone (He *et al*. 2017). WingNet was trained as an instance segmentation algorithm that identified each forewing and hindwing in the photography, masking and cropping these wings to individual image files for downstream analysis. We used a training set of 70 images (validation set, 15 images) across 1,000 epochs with a base learning rate of 0.00025. WingNet also made use of Detectron2 with a Mask-RCNN-50FPN 3x backbone (He *et al*. 2017). Loss and accuracy metrics for both CNNs can be found in Supplemental Figures S1 and S2. Melanism was extracted using the L*a*b* color value for all pixels in the wing. For this, we considered a L* value of below 10, and a* value of between -10 to 10, and a b* value between -10 to 10 as our threshold. Dark pixels falling within this threshold were classified as melanism. We validated our size measurements by ground-truthing our manual annotations with those obtained from the computer vision approach. Pearson correlation coefficients between forewing measurements (R = 0.99) and hindwing measurements (R = 0.98) were high and thus we considered our computer vision approach valid for extracting features from the images.

### Supplemental Material S2: Spatiotemporal Model Design and Selection

The random-effects structure of our model is based on a covariance matrix containing each specimen that reflected a spatiotemporal covariance structure. Calculating this matrix was essential as we lacked genetic data to construct a true phylogeny, which would typically be used to account for non-independence in this type of analysis. To calculate this matrix, we first took the distance between each photograph’s latitude and longitude coordinates. We then converted this spatial distance matrix to a spatial covariance matrix using an exponential decay function. This function was computed by:

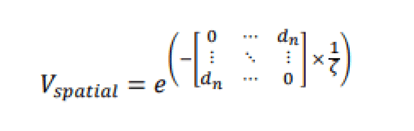

where a spatial distance matrix is multiplied by a dependency or tuning parameter, (zeta). The spatial tuning parameter,, was manually adjusted to explore scenarios where isolation-by-distance was assumed to be strong (4), slightly strong (2), moderate (1), weak (0.5), and very weak (0.25). Exponential covariance functions are widely used in spatial modeling (Ver Hoef *et al*. 2018). Secondly, the temporal component of this dependence structure was defined by a temporal autoregressive function (which we keep constant across all parameterizations of):

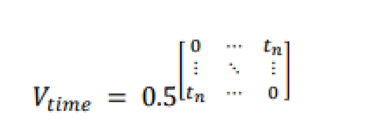

Thus, the spatiotemporal covariance between observations in our dataset is defined as a product of the spatial covariance matrix,, and the temporal covariance matrix,. We computed the full spatiotemporal covariance matrix using the package “SpatialTools” in R (French 2018).

We used the spatiotemporal covariance structure to inform the relationship between random intercepts. We calculated the variance partition coefficient (VPC) in all of our models. This metric is also sometimes called an Intraclass Correlation Coefficient (ICC). Note that in a phylogenetic context, VPC is synonymous with Pagel’s (Pagel 1999); however, we refrain from using that terminology here since our conceptualization of the relationship between individuals was calculated under the isolation-by-distance assumption rather than a phylogeny *sensu stricto*. We considered this assumption fair given the species’ largely alpine habitat and avoidance of navigating closed-canopy terrain. Further, since we varied the strength of this assumption, we were able to examine how the tuning parameter maximized the predictive ability of our models.

The AIC, log-likelihood, and VPC metrics for our models are shown in the Supplemental Table S2.2 and S2.3 below. Models were selected based on the lowest AIC score.

**Supplemental Table S2.2.** AIC scores and log-likelihoods for each model when the spatial-dependence parameter, (, was tuned to varying strengths. The top candidate models chosen for visualization are underlined, and the cell is highlighted in yellow.

| $\zeta$<br>Value | Female<br>Forewings | | Female<br>Hindwings | | Male<br>Forewings | | Male<br>Hindwings | |
| --- | --- | --- | --- | --- | --- | --- | --- | --- |
|  | AIC | LogLik | AIC | LogLik | AIC | LogLik | AIC | LogLik |
| 0.25 | <u>1281.568</u> | <u>-653.659</u> | 1154.473 | -601.307 | <u>3541.036</u> | <u>-1805.867</u> | 3054.125 | -1601.055 |
| 0.5 | 1282.025 | -654.091 | 1153.611 | -600.224 | 3542.442 | -1086.906 | <u>3053.984</u> | <u>-1600.849</u> |
| 1 | 1282.999 | -654.823 | 1152.857 | -598.956 | 3546.095 | -1808.755 | 3055.449 | -1601.563 |
| 2 | 1283.834 | -655.542 | <u>1152.670</u> | <u>-598.337</u> | 3552.004 | -1811.672 | 3059.274 | -1603.904 |
| 4 | 1283.923 | -656.014 | 1153.151 | -598.856 | 3559.197 | -1815.624 | 3064.965 | -1607.369 |

**Supplemental Table S2.3.** Variance partition coefficients (VPCs) for each model when the spatial-dependence parameter, (, was tuned to varying strengths. The top candidate models from Supplemental Table S2.2, chosen for visualization, are underlined, and the cell is highlighted in yellow.

| $\zeta$<br>Value | Female<br>Forewings | | Female<br>Hindwings | | Male<br>Forewings | | Male<br>Hindwings | |
| --- | --- | --- | --- | --- | --- | --- | --- | --- |
|  | VPC |  | VPC |  | VPC |  | VPC |  |
| 0.25 | <u>0.320</u> |  | 0.511 |  | <u>0.343</u> |  | 0.631 |  |
| 0.5 | 0.329 |  | 0.511 |  | 0.353 |  | <u>0.642</u> |  |
| 1 | 0.342 |  | 0.510 |  | 0.365 |  | 0.658 |  |
| 2 | 0.359 |  | <u>0.519</u> |  | 0.381 |  | 0.684 |  |
| 4 | 0.383 |  | 0.547 |  | 0.412 |  | 0.720 |  |

**Supplemental Figure S1.**
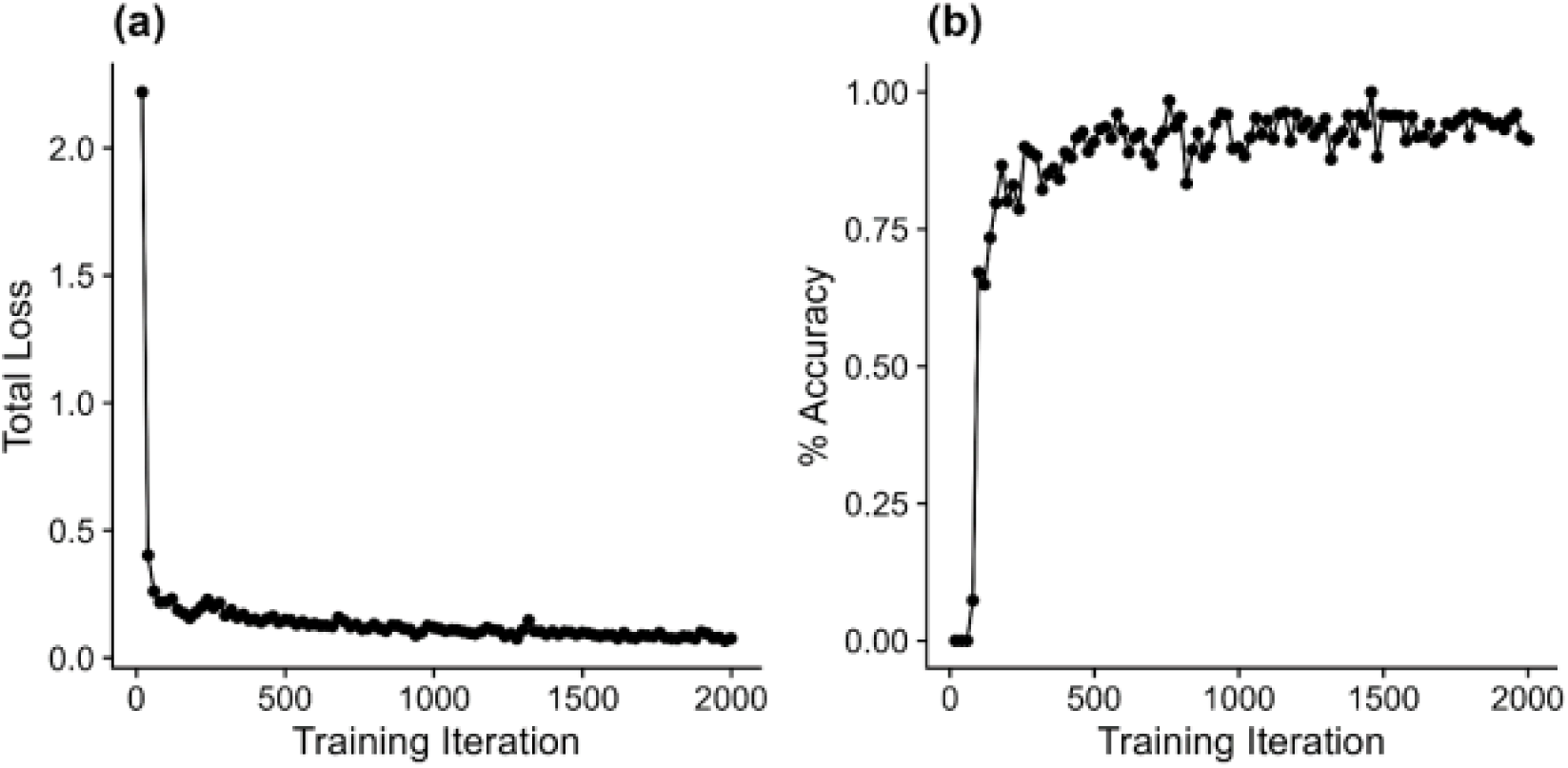
(a) Loss and (b) accuracy metrics for ScaleNet indicating the appropriate training and performance of the neural network.

**Supplemental Figure S2.**
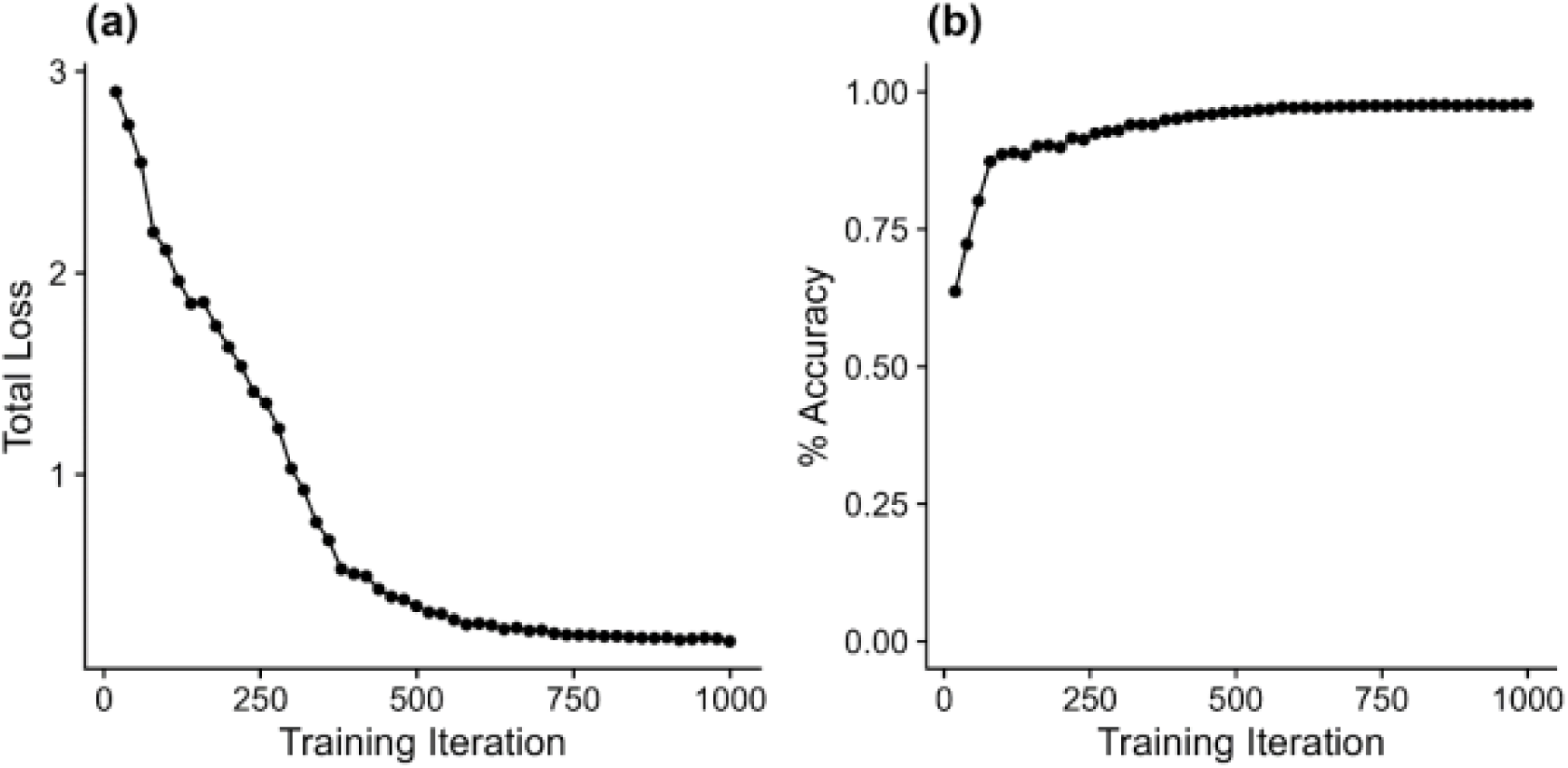
(a) Loss and (b) accuracy metrics for WingNet, indicating the appropriate training and performance of the neural network.

**Supplemental Figure S3.**
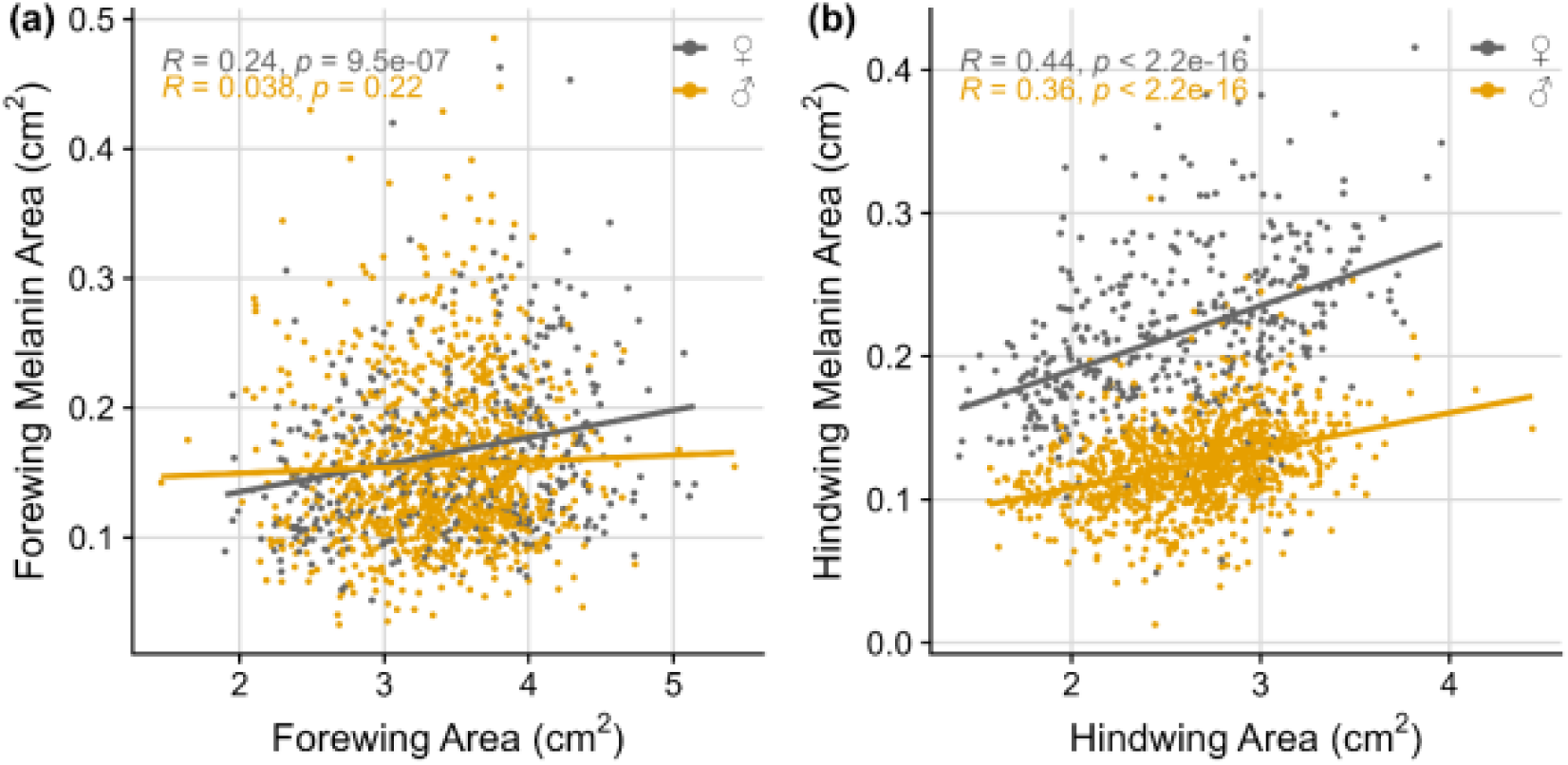
The correlation between wing area and area of melanized wing on (a) forewings and (B) hindwings for males and females.

**Supplemental Figure S4.**
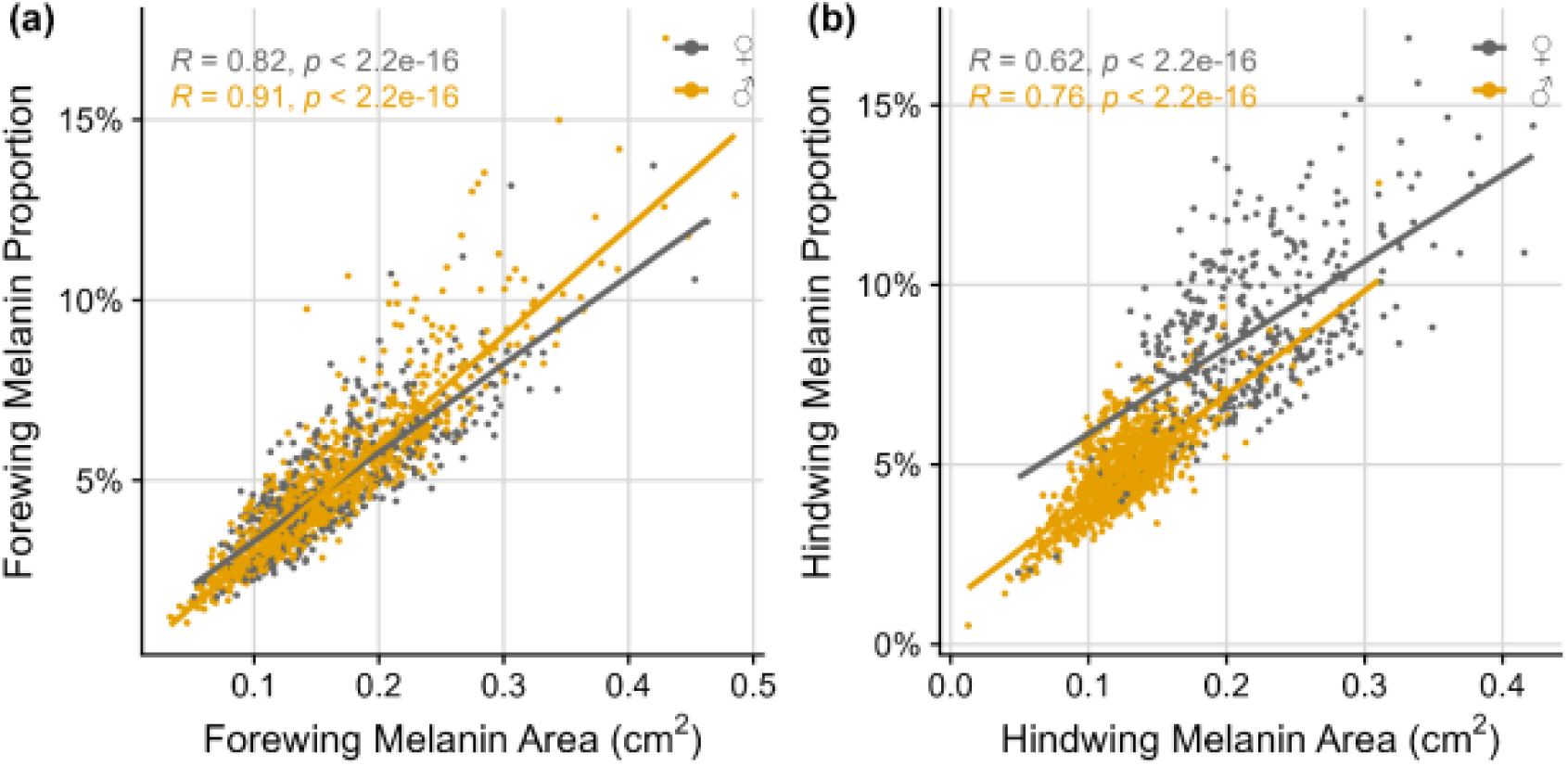
The correlation between the area of wing melanism and the wing melanization proportion on (a) forewings and (b) hindwings for males and females.

**Supplemental Figure S5.**
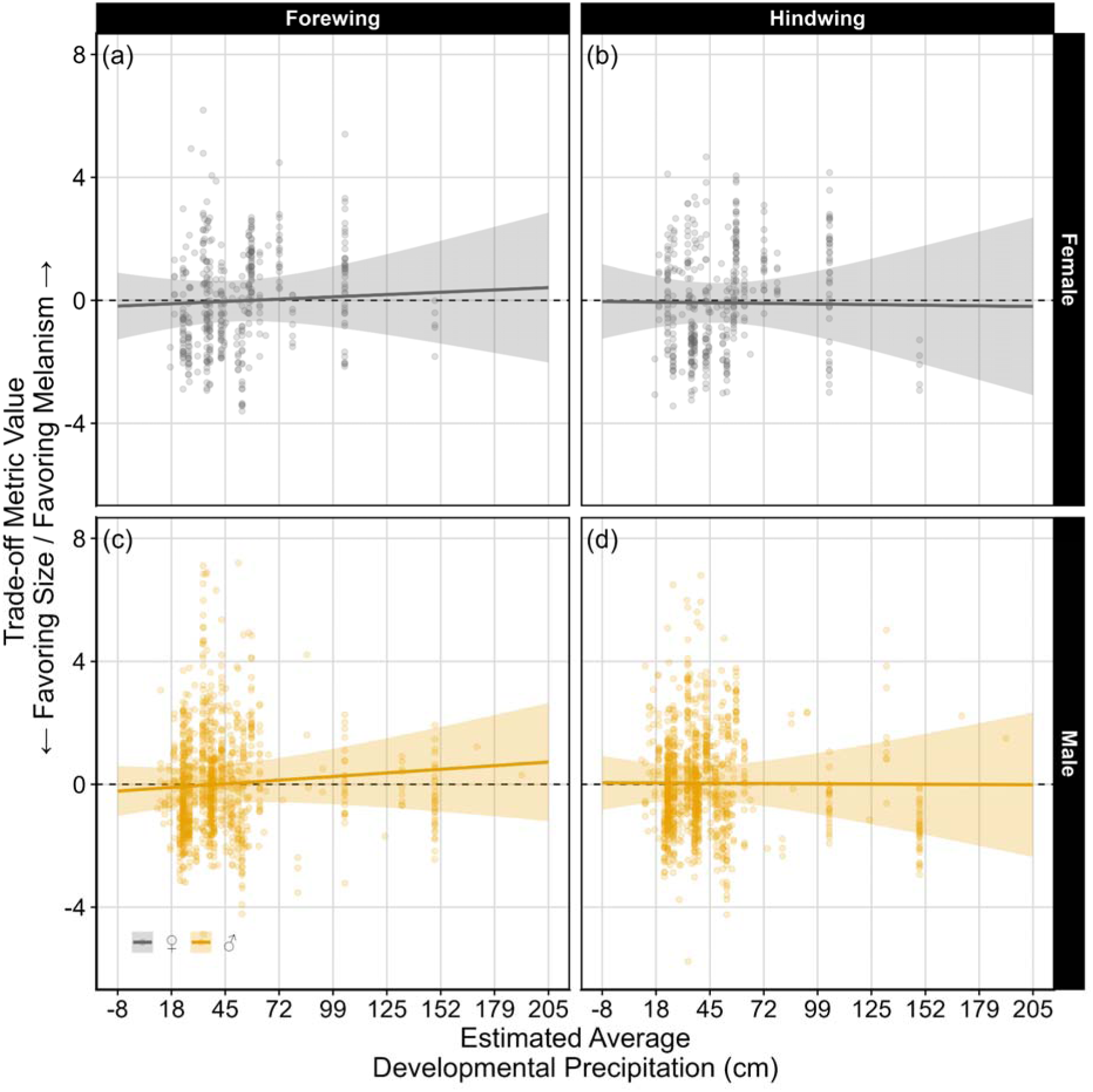
Partially-dependent effect plots of inferred average developmental precipitation on (a) female forewing, (b) female hindwing, (c) male forewing, and (d) male hindwing size-melanism trade-offs. Positive values indicate favoring melanism, while negative values indicate favoring size. The relationships between precipitation and the trade-off metric are non-significant for all sexes and wings (see Table 1 for model statistics).

